# Eosinophilic esophagitis-associated epithelial remodeling may limit esophageal carcinogenesis

**DOI:** 10.1101/2022.11.28.517589

**Authors:** Annie D. Fuller, Adam L. Karami, Mohammad Faujul Kabir, Alena Klochkova, Jazmyne L. Jackson, Anbin Mu, Yinfei Tan, Andres Klein-Szanto, Kelly A. Whelan

## Abstract

Under homeostatic conditions, esophageal epithelium displays a proliferation/differentiation gradient that is generated as proliferative basal cells give rise to suprabasal cells then terminally differentiated superficial cells. This proliferation/differentiation gradient is perturbed in esophageal pathologies both benign and malignant. Esophageal cancer is among the deadliest forms of human malignancy with 5-year survival rates of <20%. Esophageal squamous cell carcinoma (ESCC) and esophageal adenocarcinoma (EAC) are the two most common subtypes of esophageal cancer. Gastroesophageal reflux disease (GERD) is a primary risk factor for EAC. Although GERD and the food allergy-mediated condition eosinophilic esophagitis (EoE) are both associated with chronic esophageal inflammation and epithelial remodeling, including basal cell hyperplasia, epidemiological evidence suggests that EoE patients do not develop esophageal malignancy. Here, we perform single cell RNA-sequencing in murine models of EoE and ESCC to delineate the effects that these two conditions have specifically upon the cellular landscape of esophageal epithelium. In mice with EoE or ESCC, we find expansion of cell populations as compared to normal esophageal epithelium. In mice with EoE, we detect expansion of 4 suprabasal populations coupled with depletion of 4 basal cell populations. By contrast, mice with ESCC display expansion of 4 basal populations as well as depletion of 3 superficial populations. We further evaluated modules of co-expressed genes in EoE- and ESCC-enriched epithelial cell clusters. Senescence, glucocorticoid receptor signaling, and granulocyte-macrophage colony-stimulating factor pathways were associated with EoE-enriched clusters while pathways associated with cell proliferation and metabolism were identified in ESCC-enriched clusters. Finally, by pairing murine models of EoE and ESCC, we demonstrate that exposure to EoE inflammation limits esophageal carcinogenesis. Our findings provide the first functional investigation of the relationship between EoE and esophageal cancer and suggest that esophageal epithelial remodeling events occurring in response to EoE inflammation may limit act to esophageal carcinogenesis which may have future implications for leveraging allergic inflammation-associated alterations in epithelial biology to prevent and/or treat esophageal cancer.

## 1 Introduction

Esophageal epithelium exhibits an exquisite proliferation/differentiation gradient under homeostatic conditions. In this stratified squamous epithelium, proliferation is confined to the basal cell layer (1, 2). As basal cells migrate outward toward the lumen, they execute a terminal differentiation program, giving rise to overlying suprabasal cells then terminally differentiated superficial cells (3). Maintenance of this proliferation/differentiation gradient, which is essential for epithelial barrier function, is perturbed in esophageal pathologies.

Esophageal squamous cell carcinoma (ESCC) is the most common form of esophageal cancer worldwide. ESCC arises via malignant transformation of esophageal epithelial cells with activation of epidermal growth factor receptor (EGFR) and cyclin D1 oncogenes and mutations in *TP53* tumor suppressor gene representing common genetic alterations (4–8). The histological progression from normal esophageal epithelium to ESCC includes stepwise progression from basal cell hyperplasia to intraepithelial neoplasia, dysplasia, then frank carcinoma. By contrast, esophageal adenocarcinoma (EAC) develops as esophageal epithelium is displaced by specialized intestinal columnar mucosa. This metaplastic condition, termed Barret’s esophagus (BE), is associated with gastroesophageal reflux disease (GERD) and represents the strongest known risk factors for EAC (9, 10). GERD is a form of esophagitis through which acid-rich refluxate from the lower gastrointestinal tract enters the esophagus, inducing both chemical injury and cytokine-mediated injury to esophageal mucosa as well as epithelial remodeling in the form of basal cell hyperplasia (11, 12).

Eosinophilic esophagitis (EoE) is a chronic allergen-induced inflammatory disorder affecting ~4 in 10,000 individuals in the United States with clinical manifestations including dysphagia, food impaction and stricture (13). Although the disease is characterized by esophageal eosinophilia and T helper (Th)2-type inflammation, changes in esophageal epithelial tissue architecture are also present in EoE patients with impaired squamous differentiation and basal cell hyperplasia contributing to barrier defects (14–17). While chronic inflammation resulting in epithelial remodeling is a hallmark of both GERD and EoE, epidemiological studies have failed to detect esophageal cancer in EoE patients (18–20). The largest of these studies retrospectively analyzed health care data for EoE, GERD, and BE patients in relation to a control cohort, and recapitulated enhanced esophageal cancer risk in both BE and GERD patients, but not those with EoE (20). These data raise possibility that EoE inflammation may exert a protective effect with regard to esophageal carcinogenesis.

Various studies have indicated negative associations between atopy and cancer risk (21–30). While emerging evidence supports immune-mediated mechanisms, including enhanced immunosurveillance and suppression of tumor-eradicating Th1 inflammation (31–34), as potential factors supporting these epidemiological findings, we postulated that alterations in the cells that give rise to tumors may also support impaired carcinogenesis in atopic individuals. To test this hypothesis, the current study explores the impact of ESCC and the food allergen-mediated disorder EoE upon the esophageal epithelial landscape using single cell RNA-Sequencing (scRNA-Seq). We focus on EoE with ESCC for the current study as (1) robust murine models of these two conditions are available; and (2) ESCC arises from direct transformation of esophageal keratinocytes. Our studies reveal that exposure to EoE inflammation drives accumulation of suprabasal populations coupled with depletion of basal populations. By contrast, ESCC induces accumulation of basal populations concomitant with depletion of superficial populations. Pathway analysis of genes displaying co-expression further indicates that epithelial remodeling in EoE is associated with senescence, glucocorticoid signaling, and granulocyte-macrophage colony-stimulating factor (GM-CSF) signaling while epithelial remodeling in ESCC is associated with cell proliferation and cell metabolism pathways. Finally, we report that that exposure to EoE inflammation limits ESCC carcinogenesis *in vivo*, providing the first functional interrogation of the relationship between these two esophageal pathologies.

## 2 Material and Methods

### Animal Experiments

All research for the current study complies with all relevant ethical regulations. All murine studies were performed in accordance with a protocol approved by Temple University IACUC (Protocol Number: 5018). All animal experiments were conducted in accordance with institutional guidelines for animal research. All mice were maintained under controlled conditions with a 12 h light/dark cycle at an appropriate temperature and humidity. C57BL/6 mice (Cat# 000664) were obtained from the Jackson Laboratory (USA) and bred for experiments. In mice, administration of the food allergen ovalbumin (OVA; A5503, Sigma-Aldrich, St. Louis, MO, USA) coupled with cutaneous challenge with the Vitamin D analog MC903 (calcipotriol; 2700, Tocris, Bristol, UK) promotes esophageal eosinophilic infiltrates and food impactions (28, 35) and oral administration of the carcinogen 4-nitroquinoline 1-oxide (4NQO; N8141, Sigma-Aldrich) induces esophageal tumors that recapitulate histological and molecular features of human ESCC (36, 37).

The following procedures were conducted for individual induction of EoE and ESCC. EoE-like inflammation was induced using the previously described MC903/OVA mouse model (35, 38) over a period of 32 days. For 12 days, ears of mice were scraped with a scalpel blade then 20 μl MC903 (10 μM dissolved in 100% ethanol) was applied to each ear followed by 10 μl OVA (10 μM in PBS). From days 15-32, mice were subjected to oral gavage with 100 μl OVA (500 mg/ml in water) every other day and provided *ad libitum* access to drinking water supplemented with OVA (15 g/L). Mice were euthanized at day 32 and esophagi were dissected for scRNA-Seq analysis. To induce ESCC, mice were administered 4NQO (100 μg/mL in 2% propylene glycol) for 16 weeks via drinking water. 4NQO was then withdrawn for a period of 8 weeks. At the end of this 24-week protocol, mice were euthanized, and esophagi were dissected for scRNA-Seq analysis. Untreated wild type C57BL/6 mice served as controls.

For experiments combining EoE and ESCC, C57BL/6 mice were randomly assigned to one of 4 treatment groups and the following procedures were conducted over a period of 28 weeks.

- **Mice in Group 1 [EoE (-) ESCC (-)]** served as a MC903-only control. As MC903 induces dermatitis, it is important to control for the effects of this agent with regard to esophageal carcinogenesis (39). Mice in group 1 were treated with MC903 only for a period of 32 days. For 12 days ears of mice were scraped with a scalpel blade then 20 μl MC903 (10 μM dissolved in 100% ethanol) was applied to each ear followed by 10 μl PBS. From days 15-32, mice were subjected to oral gavage with 100 μl water every other day. At day 33, mice were provided drinking water with 2% propylene glycol for 16 weeks. Mice were then administered normal drinking water for 8 weeks. For the final 24 weeks of the experiment mice, were subjected to oral gavage with water every other day for the last week of each month.
- **Mice in Group 2 [EoE (+) ESCC (-)]** served to assess the long-term effects of EoE exposure with regard to esophageal epithelial alterations. Mice in group 2 were treated with MC903 and OVA for a period of 32 days. For 12 days, ears of mice were scraped with a scalpel blade then 20 μl MC903 (10 μM dissolved in 100% ethanol) was applied to each ear followed by 10 μl OVA (10 μM in PBS). From days 15-32, mice were subjected to oral gavage with 100 μl OVA (500 mg/ml in water) every other day and provided *ad libitum* access to drinking water supplemented with OVA (15 g/L). At day 33, mice were provided drinking water with 2% propylene glycol for 16 weeks. Mice were then administered normal drinking water for 8 weeks. During the 24-week period following EoE induction, mice were subjected to oral gavage with 100 μl OVA (500 mg/ml in water) every other day and provided *ad libitum* access to drinking water supplemented with OVA (15 g/L) for the last week of each month.
- **Mice in group 3 [EoE (-) ESCC (+)]** served to assess the effect of 4NQO alone on esophageal carcinogenesis. Mice in group 3 were treated with MC903 only for a period of 32 days. For 12 days, ears of mice were scraped with a scalpel blade then 20 μl MC903 (10 μM dissolved in 100% ethanol) was applied to each ear followed by 10 μl PBS. From days 15-32, mice were subjected to oral gavage with 100 μl water every other day. At day 33, mice were provided drinking water with 4NQO (100 μg/mL in 2% propylene glycol) for 16 weeks. Mice were then administered normal drinking water for 8 weeks. For the final 24 weeks of the experiment mice, were subjected to oral gavage with water every other day for the last week of each month.
- **Mice in Group 4 [EoE (+) ESCC (+)]** served to assess the effects of EoE exposure on esophageal carcinogenesis. Mice in group 4 were treated with MC903 and OVA for a period of 32 days. For 12 days, ears of mice were scraped with a scalpel blade then 20 μl MC903 (10 μM dissolved in 100% ethanol) was applied to each ear followed by 10 μl OVA (10 in PBS). From days 15-32, mice were subjected to oral gavage with 100 μl OVA (500 mg/ml in water) every other day and provided *ad libitum* access to drinking water supplemented with OVA (15 g/L). At day 33, mice were provided drinking water with 4NQO (100 μg/mL in 2% propylene glycol) for 16 weeks. Mice were then administered normal drinking water for 8 weeks. During the 24-week period following EoE induction, mice were subjected to oral gavage with 100 μl OVA (500 mg/ml in water) every other day and provided *ad libitum* access to drinking water supplemented with OVA (15 g/L) for the last week of each month.

### scRNA library preparation and sequencing

Single cell droplets were generated with inDrop according to manufacturer’s protocols. 5000–7000 cells were collected to make cDNA at the single cell level. Full-length cDNA with Unique Molecular Identifiers (UMI) was synthesized via reverse-transcription in the droplet. After PCR amplification and purification, cDNA was fragmented to ~270 bp and the Illumina adapters with index were ligated to fragmented cDNA.

### Deconvolution of scRNA-Seq Reads

FASTQ files from the sequencing run were downloaded from Illumina’s BaseSpace sequence hub. To resolve the mapping of cellular barcodes and Unique Molecular Identifiers (UMIs), UMI-tools (v1.0.0) was used to whitelist and extract the barcodes. Likely barcodes were found using the whitelist function of UMI-tools, which searches for the inDrop regular expression pattern of “ (?P<cell_1>.(7))(?P<discard_1>GAGTGATTGCTTGTGACGCCTT){s<=2}(?P<cell_2>.{8})(?P<umi_1>.{6})T(42).*” in the R2 of the read pairs. Using a whitelist of likely barcodes, the extract function relocated both the cell barcode and the unique molecular identifier found in the same read to the read name in the FASTQ files. The extraction retains the information of unique cellular barcodes while enabling correct read mapping of genes without the attached cell and transcript identifiers. Reads were then mapped using the aligner STAR (v2.7.3). First, the murine genome index was made using the M23 GRCm38 genomic sequence from GENCODE. Using the index, all read pairs were aligned to the genome for an output of BAM files. To limit the variance of amplification on the length of transcripts, the UMI-tools dedup tool was used to deduplicate the reads, which deduplicates UMIs that match to a single transcript. The resulting deduplicated alignments were then mapped to the murine gene transfer file (GTF) from GENCODE using featureCounts to count the number of reads mapping to each gene in the genome. Finally, the UMI-Tools count function was used to summarize the gene counts in each cell in each sample to give an output of a matrix with gene names and cell barcodes.

### Data Filtration, Initial Clustering, and Exclusion of Non-Epithelial Cells

The matrices for each sample were imported and transformed into Seurat (v3.2.2) objects for further processing. Genes expressed in 3 or fewer cells are excluded from analysis. Doublet and dead cell removal were done and based on the count distribution of each sample. To equalize cell counts between conditions, 19,041 total cells sequenced were downsampled to 1,500 cells per condition (4,500 cells total).

Using Monocle3 on R (v1.0.0), a gene expression matrix (GEM) was generated from Seurat objects and used to create a cell data set (CDS) object. The resulting dataset was then scaled and centered for dimensionality reduction. PCA was used for initial dimensionality reduction and later for clustering, resulting in 30 principal components. The PCA principal components were then used as input to the Uniform Manifold Approximation and Projection (UMAP) dimensionality reduction procedure for visualization, using 30 neighbors for local neighborhood approximation and embedding into 2 components for visualization. Cell population clusters were developed using Monocle3. 13 distinct cellular populations were found, each with a shared transcriptomic profile across different samples.

### Gene Module Discovery

To determine transcriptomic programs prevalent in the different subsections of the epithelial dataset and across pathological conditions, we used Monocle3’s gene module functionality. Briefly, Monocle3 performs UMAP on genes with cells as features to find distinct groups of genes locally co-expressed in the spatio-pseudotemporal trajectory inferred within the cellular UMAP space. 20 such gene modules were found. Modules of pathological interest were imported into QIAGEN Ingenuity Pathway Analysis (IPA) for core analysis.

### Cell Cycle Phase Prediction and Pseudotemporal Trajectory Inference

Pre-processing for pseudotemporal trajectory inference included estimation of size factors and dispersion and choosing genes to be used for ordering the cells in pseudotime and clustering according to the “dpFeature” procedure on Monocle3. The root node was isolated at the crux of the G1/G0 and G2M/S phases of the cell cycle as determined by the Seurat function “CellCycleScoring”. Unsupervised reverse graph embedding delineated branch points and pseudotime trajectories on Monocle3.

### Statistical Analysis

Unpaired Student’s t-test, two-way ANOVA followed by Tukey’s multiple comparisons, Wilcoxon signed-rank test, and Fisher’s exact test (as indicated in figure legends) were used for statistical evaluation of data. p<0.05 was used as the threshold for statistical significance. Statistical analysis was performed using GraphPad Prism (GraphPad Software, La Jolla, CA, USA).

## 3 Results

### Identification and characterization of esophageal epithelial cell populations along the basal/superficial cell axis in mice with EoE or ESCC

To define how EoE and ESCC influence the cellular landscape of esophageal epithelium, we performed scRNA-Seq on mice treated with MC903/OVA to induce EoE (n=3), 4NQO to induce ESCC (n=4), or untreated controls (n=12). Epithelium was peeled from dissected esophagi of all mice and subjected to scRNA-Seq using the inDrop platform (**Figure 1A**). The resulting dataset consisted of 4,500 cells across the three groups that were then subjected to unsupervised dimensionality reduction and visualized by UMAP (**Figure 1B**). Monocle3-based clustering within the dataset further revealed 13 epithelial cell populations and the top 5 most upregulated genes in each population (**Figure 1C, D**). To establish the identity of these populations, we employed *Krt5* (encoding Cytokeratin 5) and *Krtdap* (encoding Keratin Differentiation-Associated Protein) as these are respective markers of basal and superficial cells (**Figure 2A, B**) (40). Relative *Krt5* vs. *Krtdap* expression for all populations was then plotted and evaluated relative to a linear slope of 1. As suprabasal cells are expected to minimally express both basal and superficial markers, cell populations within our data set with <0.5 log_10_ normalized expression value for both *Krt5* and *Krtdap* were defined as suprabasal. Among populations with >0.5 log_10_ normalized expression value for both *Krt5* and *Krtdap*, those falling below the slope line were defined as basal and those falling above were defined as superficial (**Figure 2C**). Using these methods, 6 basal, 5 suprabasal, and 2 superficial populations were identified within the dataset (**Figure 2C, D**). Mapping of additional established markers of basal and superficial cells in esophageal epithelium supported these classifications (**Figure 2E**).

**Figure 1.**
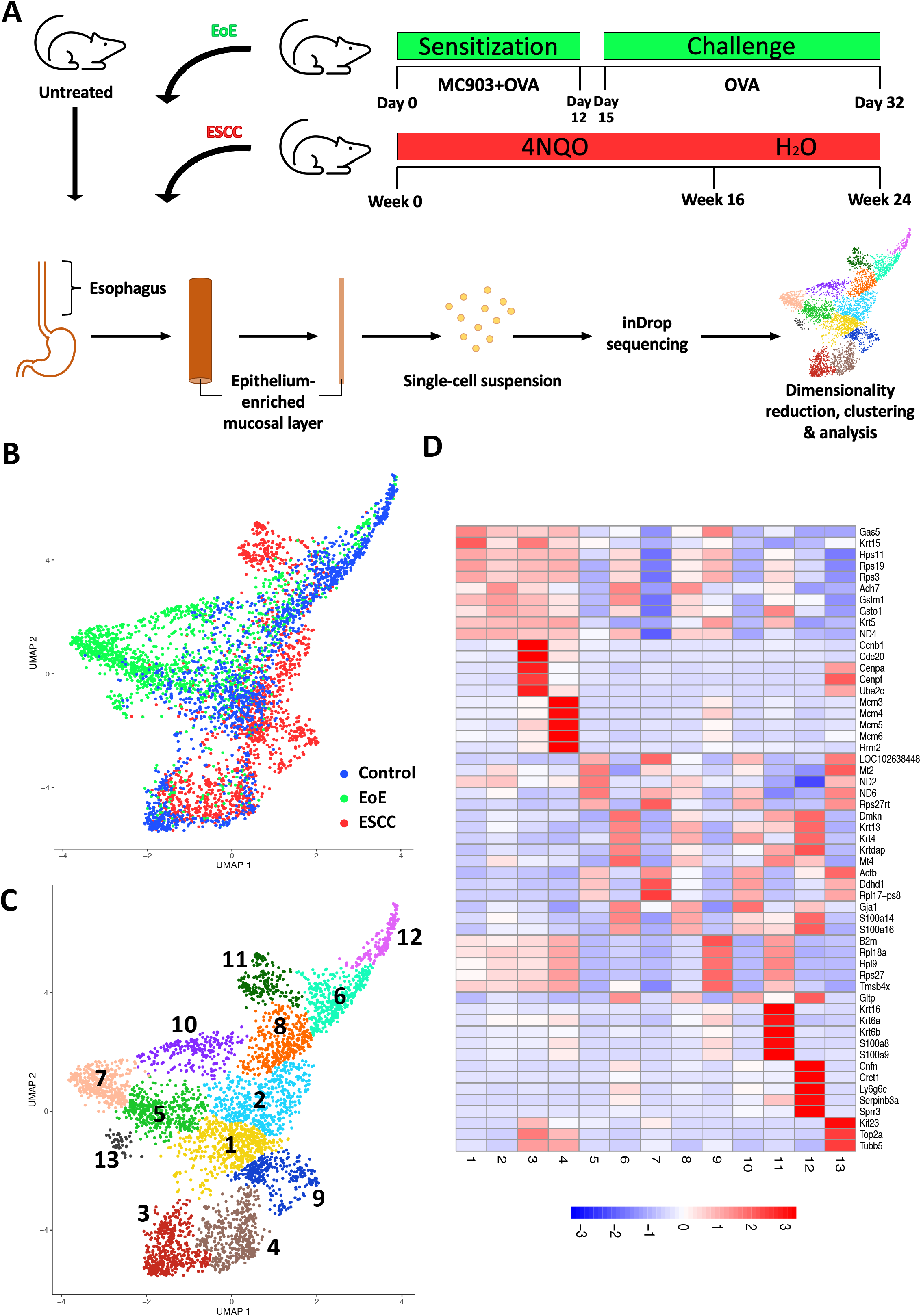
Single cell profiling of esophageal epithelium from mice treated with eosinophilic esophagitis (EoE) or esophageal squamous cell carcinoma (ESCC). **(A)** Schematic of experimental approach. MC903 and Ovalbumin (OVA) were used to induce EoE over 32 days in C57B6 mice (n=4). During the 12-day sensitization period, mice were treated epicutaneously with MC903 and OVA. Challenge with OVA in drinking water and via gavage (3x/week) was conducted from day 15 through day 32. ESCC was induced in C57B6 mice (n=3) using the chemical carcinogen 4-nitroquinoline-1-oxide (4NQO). 4NQO was administered via drinking water for 16 weeks followed by an 8 week wash out period. Untreated C57B6 mice (n=12) served as controls. In each mouse, esophageal epithelium-enriched mucosal layer was peeled from underlying muscle then enzymatically digested to generate a single cell suspension that was subjected to single cell RNA-Sequencing using the inDrop platform. **(B, C)** Uniform Manifold Approximation and Projection plot (UMAP) shows distribution of all cells from the single cell RNA-Sequencing dataset by condition with blue identifying cells from control mice, green identifying cells from mice with EoE, and red identifying cells from mice with ESCC in **B** or by distinct cell populations identified using the R-based program Monocle3 in **C**. **(D)** Heat map shows z-score scaled normalized expression of the 5 most upregulated genes in each population across all clusters. Red represents upregulation and blue represents downregulation.

**Figure 2.**
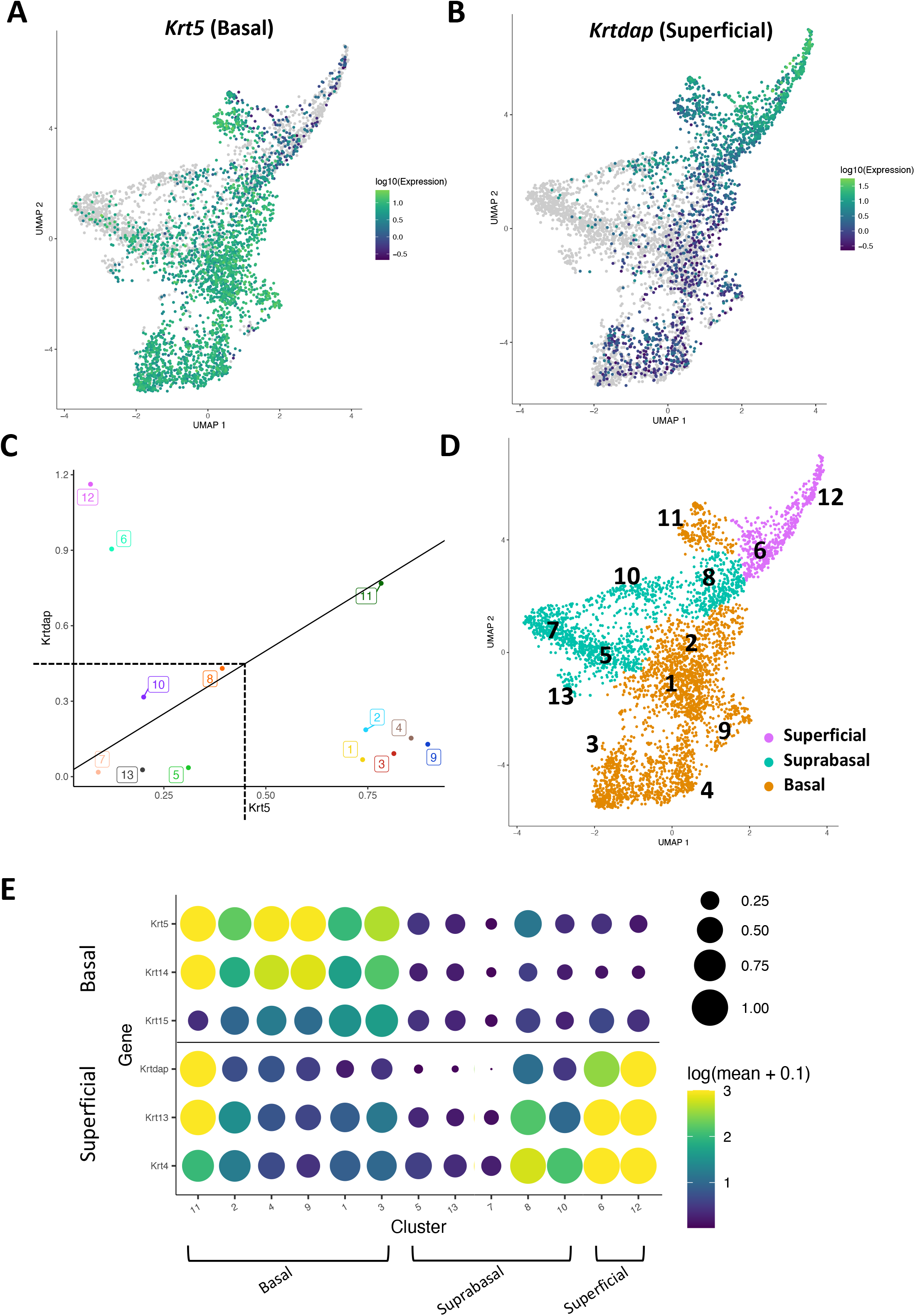
Identification of basal, suprabasal, and superficial cells across dataset. **(A, B)** Normalized log_10_ expression gradients of Keratin 5 (*Krt5*), an established marker of basal esophageal epithelial cells in **A**, or Keratinocyte Differentiation-Associated Protein (*Krtdap*), an established marker of differentiated esophageal epithelial cells in **B** are shown across the single cell RNA-Sequencing dataset. Green indicates enrichment while purple indicates inhibition. **(C)** Normalized log_10_ expression of *Krt5* vs. normalized log_10_ expression of *Krtdap* expression was plotted for all populations identified in the single cell RNA-Sequencing dataset. Populations below a linear slope of 1 were defined as basal and those above this slope line line were identified as superficial. Clusters with expression below normalized log_10_ expression of 0.5 for both markers were defined as suprabasal. **(D)** Basal (orange), suprabasal (teal), and superficial (pink) clusters identified on a Uniform Manifold Approximation and Projection plot (UMAP) of the entire single cell RNA-Sequencing dataset. **(E)** Cluster-average expression z-scores of putative basal and differentiated markers are shown for each population in the single cell RNA-Sequencing dataset. Circle size reflects percentage of cells with non-zero expression level for indicated genes. Color intensity reflects average expression level across all cells within each cluster with yellow indicating enrichment and purple indicating inhibition.

### Effects of EoE and ESCC on cell populations and trajectories in esophageal epithelium

Evaluation of our dataset separated by treatment group revealed that mice with EoE and ESCC both displayed expansion and depletion of cell populations (**Figure 3A, B**). Expansion of populations 5, 7, 10, and 13 and depletion of populations 2, 3, 4, and 9 was unique to mice with EoE. Expansion of populations 2, 4, 9, and 11 and depletion of populations 5, 8, and 12 was unique to mice with ESCC (**Figure 3A**). In mice with EoE, enriched cell populations were exclusively suprabasal (**Figure 4A**) and depleted populations were exclusively basal. By contrast mice ESCC exhibited expansion of basal cells (**Figure 4A**) and depleted populations were either superficial (i.e. population 12) or suprabasal populations (i.e. populations 5 and 8).

**Figure 3.**
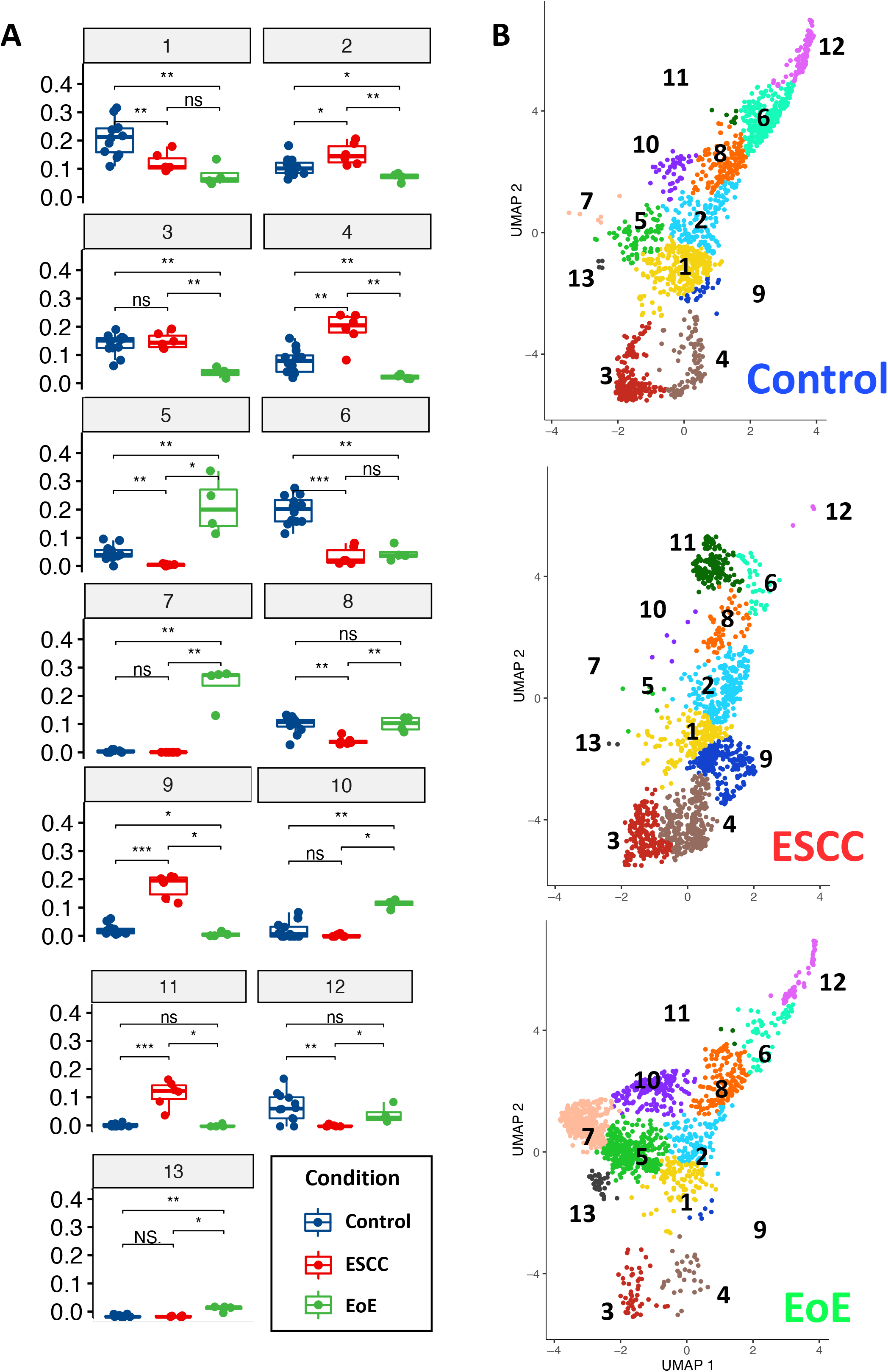
Effects of eosinophilic esophagitis (EoE) and esophageal squamous cell carcinoma (ESCC) on representation of esophageal epithelial cell populations. **(A)** Proportion of each cell as a fraction of all cells in the single cell RNA-Sequencing dataset with blue indicating control mice, red indicating mice with ESCC, and green indicating mice with EoE. Each individual scatter point represents proportion indicated, box indicates quartiles, whiskers indicate minima and maxima. Mean is indicated by line striking through box. *, p<0.05; **, p<0.01; ***, p<0.001 by Wilcoxon signed-ranked test without adjustment for multiple comparisons. **(B)** Uniform Manifold Approximation and Projection plot (UMAP) showing cell populations identified across entire single cell RNA-Sequencing dataset separated by experimental condition.

**Figure 4.**
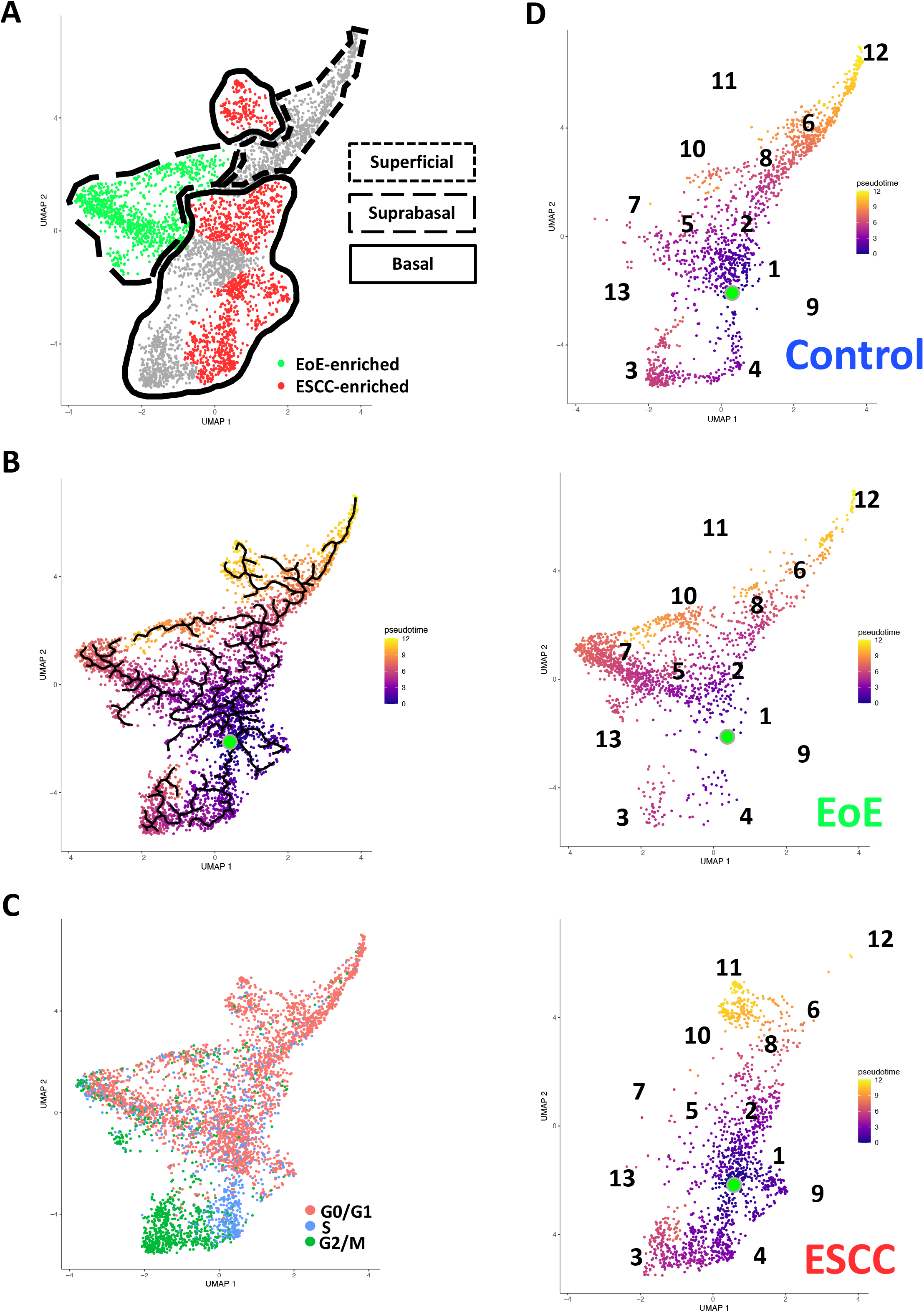
Effects of eosinophilic esophagitis (EoE) and esophageal squamous cell carcinoma (ESCC) on esophageal epithelial cell fate. **(A)** Uniform Manifold Approximation and Projection plot (UMAP) of entire single cell RNA-Sequencing dataset with cell populations found to be significantly enriched in mice with EoE or mice with ESCC colored green or red, respectively. Basal cells are enclosed in solid lines, suprabasal cells are enclosed in dashed lines, and superficial cells are enclosed in lines made of square dots. **(B)** Monocle3 UMAP visualization of all cells in single cell RNA-Sequencing dataset. Each cell is colored by its inferred pseudotime value with dark purple representing the earliest cells and bright yellow representing the latest cells in the trajectory. Green dot indicates supervised pseudotime root. Black lines represent putative cell fate trajectories. (**C**) Expression of genes associated with each phase of the cell cycle were labeled on UMAP. **(D)** Pseudotime projections of cells from untreated controls or mice with EoE or ESCC. Identification numbers show the location of individual cell populations in the pseudotime projections.

To investigate how EoE and ESCC influence cell fate in esophageal epithelial cells, we continued to perform pseudotime analysis (**Figure 4B**) coupled with evaluation of cell cycle-associated gene expression (**Figure 4C**). Based upon our prior characterization of the cellular landscape of esophageal epithelium (40), the pseudotime root was set at the G0/G1-enriched basal population immediately preceding the S phase-enriched basal population. In control mice, we detected two predominant trajectories: a cycling basal cell trajectory and a basal-suprabasal-superficial trajectory (**Figure 4B, D; Figure 5**), consistent with our previous studies (40). Pseudotime analysis further suggests that EoE-enriched suprabasal cells arise from terminal trajectories that are present in esophageal epithelium of control animals, but to a limited extent (**Figure 4B, D; Figure 5**). EoE-enriched trajectory 1 branches directly from basal cells to give rise to EoE-enriched suprabasal populations 5, 7, and 13, while EoE-enriched trajectory 2 branches from the suprabasal pool to give rise to EoE-enriched suprabasal population 10 (**Figure 4B, D; Figure 5**). Basal populations 2 and 4 are prevalent in the basal-suprabasal trajectory in control mice and are further expanded in mice with ESCC (**Figure 4B, D**). By contrast, basal populations 9 and 11 are two terminal basal cell fates that are minimally represented in control mice but are highly enriched in mice with ESCC, representing ESCC-enriched trajectories 1 and 2 (**Figure 4B, D; Figure 5**). In ESCC-enriched trajectory 1, basal population 9 branches directly from basal cells found in the normal epithelium (**Figure 4B, D; Figure 5**). In ESCC-enriched trajectory 2, basal population 11 branches off from the suprabasal cell pool common to mice in all experimental groups (**Figure 4B, D; Figure 5**), representing an aberration from the normal basal-suprabasal-superficial trajectory in esophageal epithelium.

**Figure 5.**
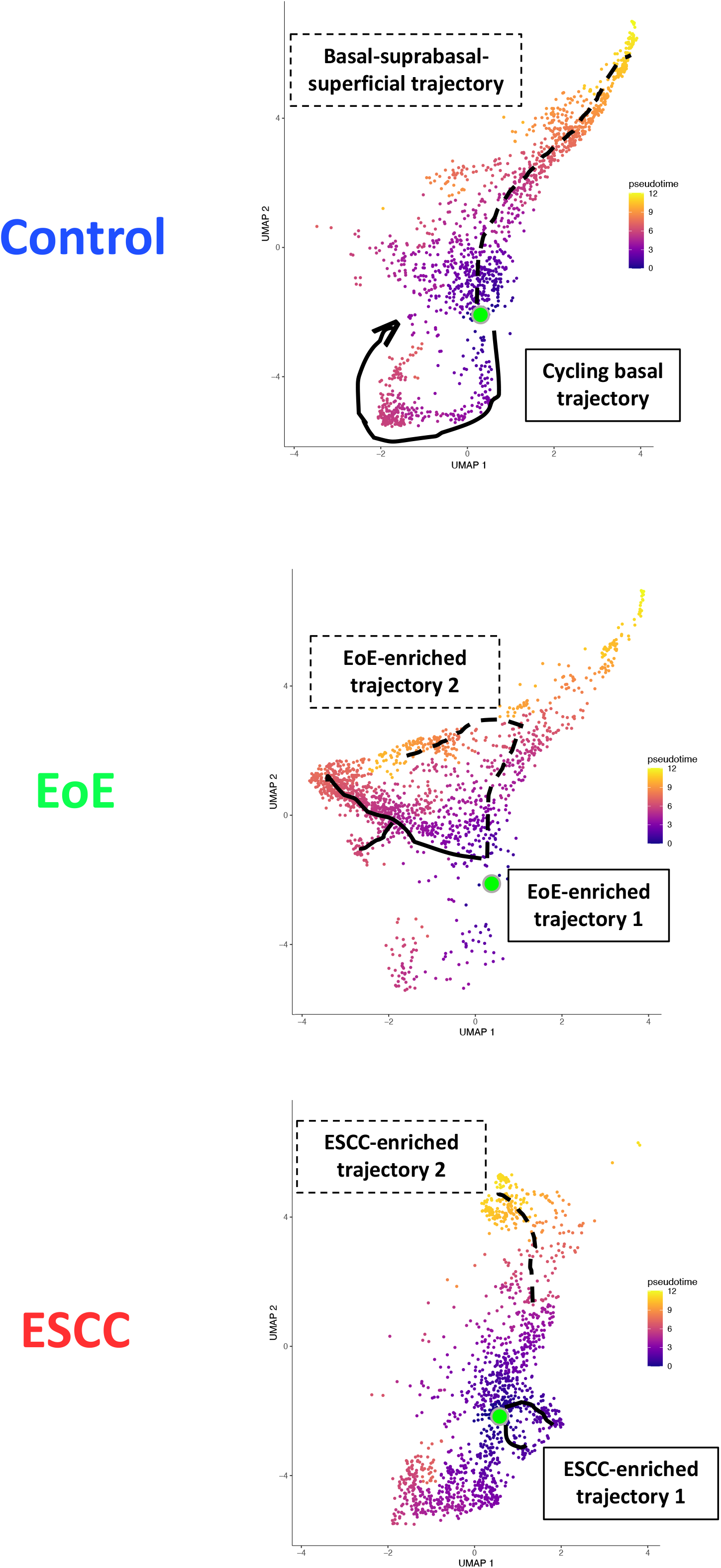
Predicted cell fate trajectories in untreated controls or mice with eosinophilic esophagitis (EoE) or esophageal squamous cell carcinoma (ESCC). Pseudotime projections of cells from untreated controls or mice with EoE or ESCC. Lines indicate predicted trajectories with trajectory names noted.

To investigate the molecular features associated with cell populations and trajectories that are enriched in mice with EoE and ESCC, co-expressed genes were grouped into modules that were differentially expressed between populations (**Figure 6**). Gene modules 3, 4, 5, and 10 were predicted to be enriched in all ESCC-specific clusters while module 12 was predicted to be highly enriched in all EoE-specific clusters (**Figure 6**). Pathway analysis revealed that genes in ESCC-enriched modules were associated with proliferation, such as EIF2 signaling, regulation of EIF4 and p70S6 kinase, and mTOR signaling in module 10 (Figure 7A; Table 1). Other pathways regulated in ESCC-enriched populations were involved in altered metabolism, such as mitochondrial dysfunction and oxidative phosphorylation in modules 3, 5 and 10 (**Figure 7A; Table 1**). Additionally, genes in module 4 were associated with molecular mechanisms of cancer (**Figure 7A; Table 1**). Whereas genes associated with proliferation were linked to ESCC-enriched clusters, EoE-enriched clusters were associated with the senescence pathway (**Figure 7B; Table 1**). Genes associated with glucocorticoid receptor signaling, which is a target for corticosteroid-based therapy in EoE patients (41), and aryl hydrocarbon receptor pathway, which has been linked to proton pump inhibitor-mediated inhibition of epithelial proliferation and IL-13 signaling (42), were also identified in EoE-enriched clusters (**Figure 7B; Table 1**). Furthermore, genes associated with granulocyte macrophage colony-stimulating factor were found in EoE-enriched clusters, consistent with the presence of eosinophils and other granulocytes in EoE patients.

**Figure 6.**
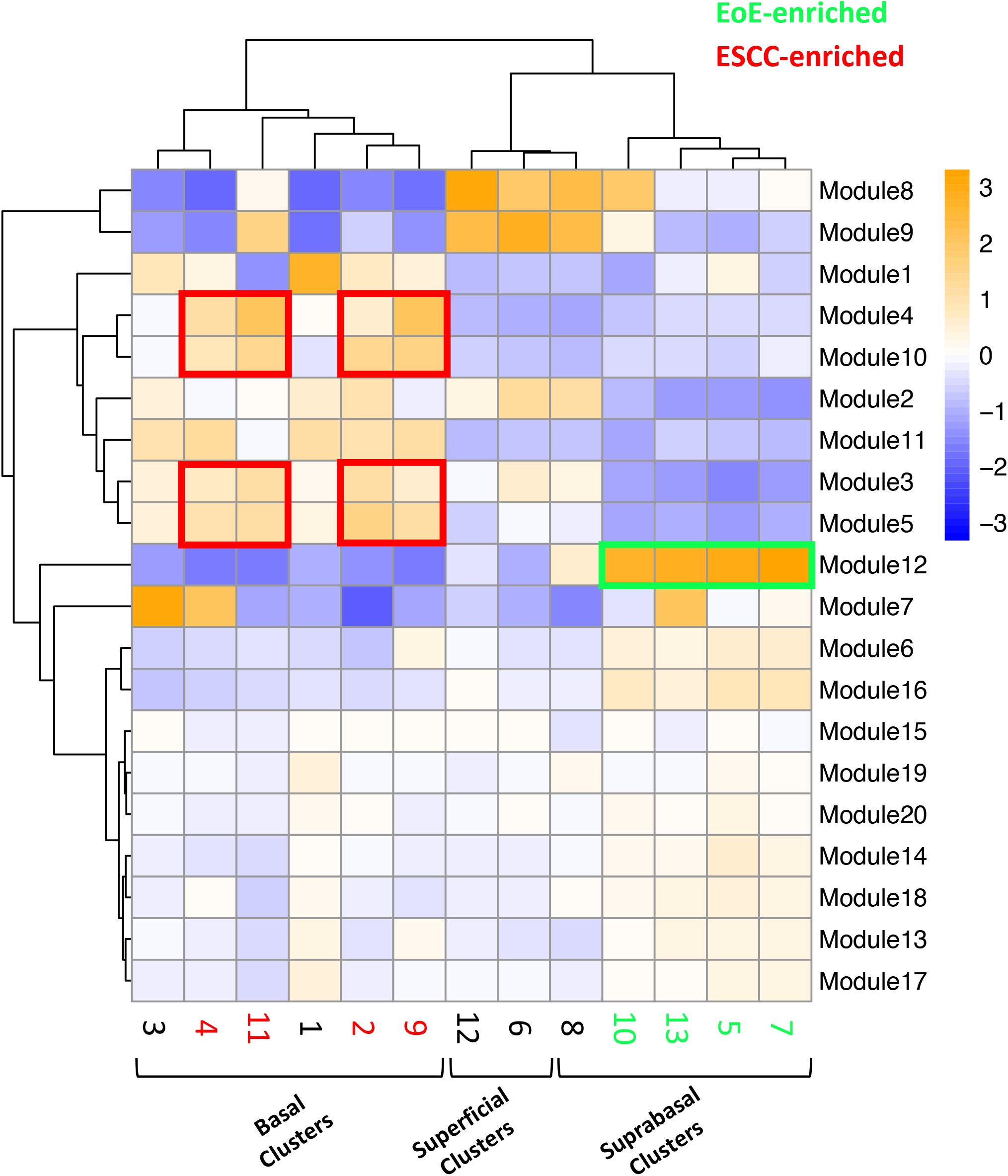
Modules of co-expressed genes and their z-score scaled normalized expression in cell populations across the single cell RNA-Sequencing dataset. A positive value (orange) indicates upregulation while a negative value (blue) indicates inhibition of the respective genes in each module. Cell populations and modules that are enriched in mice with eosinophilic esophagitis (EoE) are highlighted in green. Cell populations and modules that are enriched in mice with esophageal squamous cell carcinoma (ESCC) are highlighted in red.

**Figure 7.**
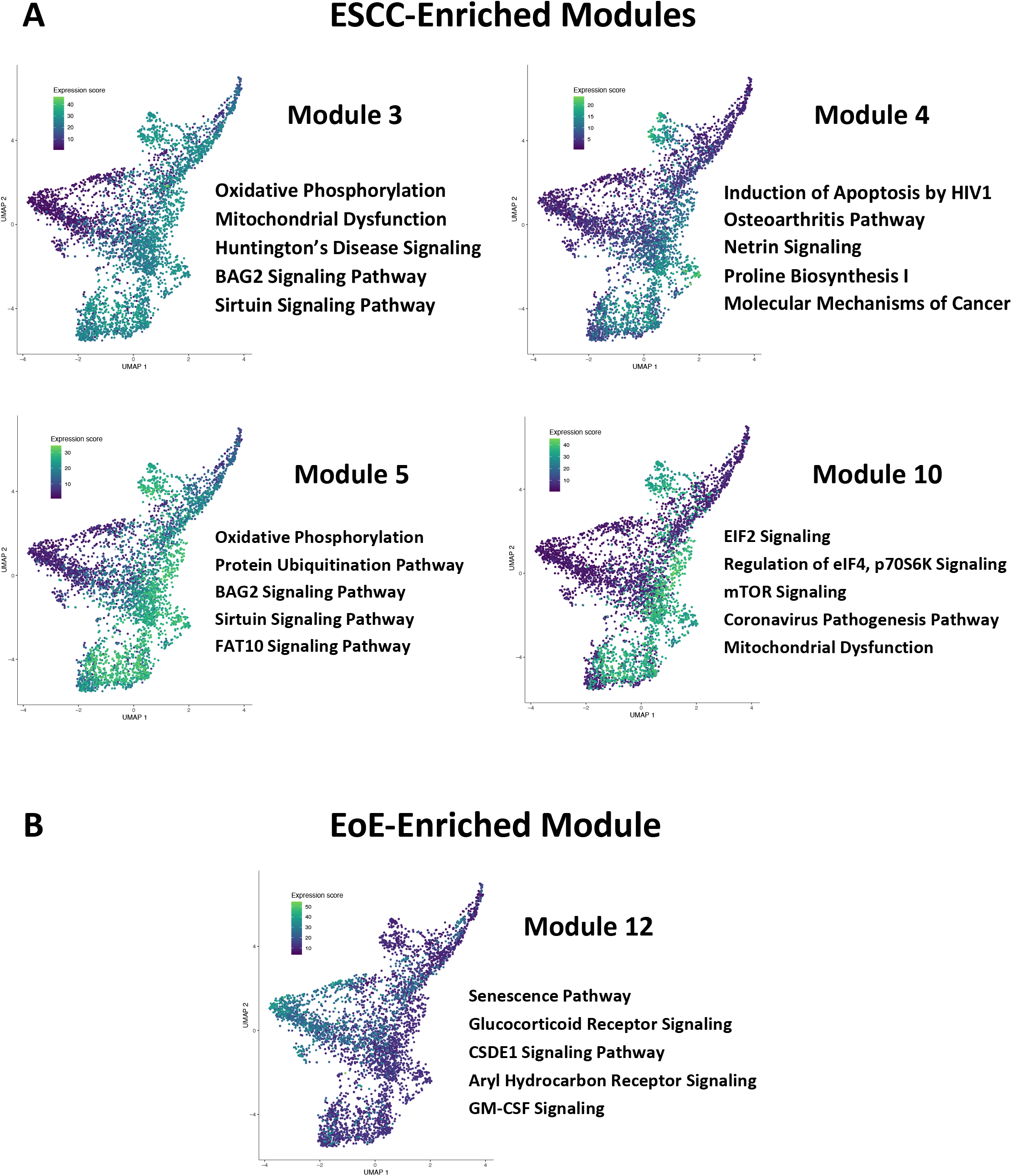
Canonical pathways associated with cell populations that are enriched in mice with esophageal squamous cell carcinoma (ESCC) or eosinophilic esophagitis (EoE). **(A, B)** Ingenuity Pathway Analysis (IPA) identified cellular processes predicted to be significantly altered in co-expressed genes in modules that are associated with populations enriched in mice with ESCC in **A** or EoE in **B.** Associated Uniform Manifold Approximation and Projection plots (UMAPs) show normalized expression levels of each module across the entire single cell RNA-Sequencing dataset with green indicating high expression and purple indicating low expression. Top 5 pathways predicted to be most significantly associated with each gene module are listed.

**Table 1.**
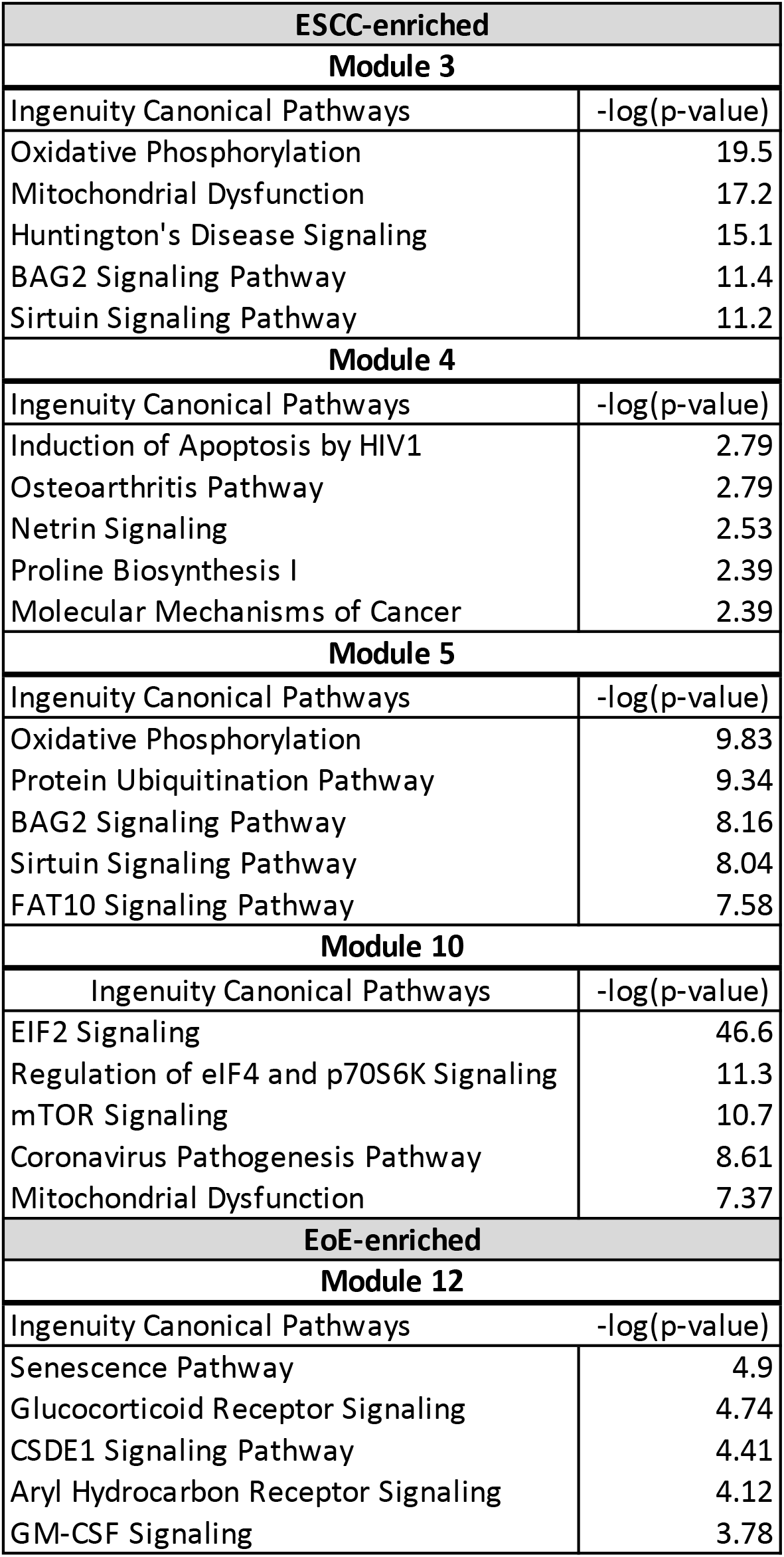
Gene modules predicted to be associated with epithelial populations that are enriched in mice with esophageal squamous cell carcinoma (ESCC) or eosinophilic esophagitis (EoE).

| **ESCC-enriched** |  |
| --- | --- |
| **Module 3** |  |
| Ingenuity Canonical Pathways | -log(p-value) |
| Oxidative Phosphorylation | 19.5 |
| Mitochondrial Dysfunction | 17.2 |
| Huntington's Disease Signaling | 15.1 |
| BAG2 Signaling Pathway | 11.4 |
| Sirtuin Signaling Pathway | 11.2 |
| **Module 4** |  |
| Ingenuity Canonical Pathways | -log(p-value) |
| Induction of Apoptosis by HIV1 | 2.79 |
| Osteoarthritis Pathway | 2.79 |
| Netrin Signaling | 2.53 |
| Proline Biosynthesis I | 2.39 |
| Molecular Mechanisms of Cancer | 2.39 |
| **Module 5** |  |
| Ingenuity Canonical Pathways | -log(p-value) |
| Oxidative Phosphorylation | 9.83 |
| Protein Ubiquitination Pathway | 9.34 |
| BAG2 Signaling Pathway | 8.16 |
| Sirtuin Signaling Pathway | 8.04 |
| FAT10 Signaling Pathway | 7.58 |
| **Module 10** |  |
| Ingenuity Canonical Pathways | -log(p-value) |
| EIF2 Signaling | 46.6 |
| Regulation of eIF4 and p70S6K Signaling | 11.3 |
| mTOR Signaling | 10.7 |
| Coronavirus Pathogenesis Pathway | 8.61 |
| Mitochondrial Dysfunction | 7.37 |
| **EoE-enriched** |  |
| **Module 12** |  |
| Ingenuity Canonical Pathways | -log(p-value) |
| Senescence Pathway | 4.9 |
| Glucocorticoid Receptor Signaling | 4.74 |
| CSDE1 Signaling Pathway | 4.41 |
| Aryl Hydrocarbon Receptor Signaling | 4.12 |
| GM-CSF Signaling | 3.78 |

### Exposure to EoE inflammation limits esophageal tumorigenesis *in vivo*

As our data indicate that EoE and ESCC drive distinct cell fate trajectories in esophageal epithelium and epidemiological data suggest that EoE patients fail to develop esophageal malignancy, we finally paired murine models of EoE and ESCC to explore the functional relationship between the two conditions (**Figure 8A**). As expected, no tumors were detected in the absence of ESCC-inducing carcinogen treatment either with [ESCC (-) EoE (+)] or without EoE [ESCC (-) EoE (-)] (**Figure 8C**). In mice treated with ESCC-inducing carcinogen, tumors were detected in 100% of mice in the absence of EoE [ESCC (+) EoE (-)] and 80% of mice in the presence of EoE [ESCC (+) EoE (+)]. Although this difference was not statistically significant, we did detect a significant decrease in tumor load in mice treated with ESCC-inducing carcinogen in the presence of EoE [ESCC (+) EoE (+)] (**Figure 8B, C**). Additionally, the spectrum of ESCC lesions was shifted in ESCC (+) EoE (+) mice in which no invasive ESCC was detected (**Figure 8D**). Furthermore, the total percentage of esophageal epithelium occupied by neoplastic lesions was significantly diminished in ESCC (+) EoE (+) mice as compared to their ESCC (+) EoE (-) counterparts (**Figure 8E**).

**Figure 8.**
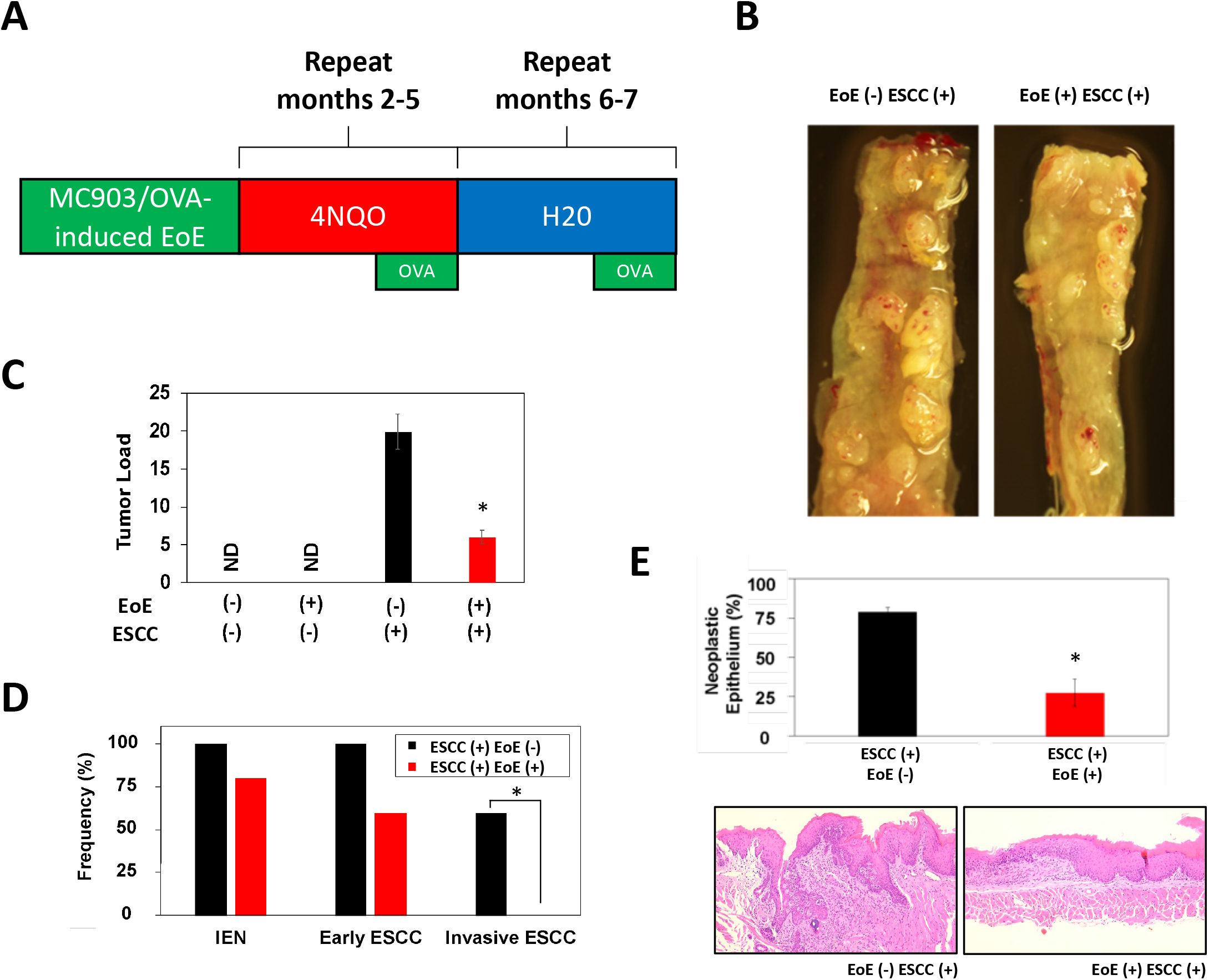
EoE inflammation limits esophageal carcinogenesis *in vivo*. **(A)** Schematic of experimental approach to combined pathology model. C57B6 mice were exposed to MC903/Ovalbumin (OVA) for one month to promote EoE, followed by exposure to the esophageal carcinogen 4-nitroquinoline 1-oxide (4NQO) to induce epithelial tumorigenesis. Esophagi were intermittently challenged with OVA to stimulate EoE-associated inflammation. Mice were either treated with vehicle controls (n=6), MC903/OVA only (n=4), 4NQO only (n=10), or MC903/OVA in combination with 4NQO (n=5). **(B)** Representative esophagi of mice treated with MC903/OVA to promote EoE and/or 4NQO to induce carcinogenesis. **(C)** Quantification of average tumor load (tumor size in mm x tumor number). ND, not detected. **(D, E)** Histological assessment was performed to quantify frequency of esophageal lesion types in **D** and percent of esophageal epithelium occupied by neoplastic lesions (squamous dysplasia or greater) with representative images shown in **E**. *, p<0.05 as calculated by unpaired student’s t-test in **C** and **E** or Fisher’s exact test in **D**.

## 4 Discussion

Esophageal epithelial remodeling has been histologically documented during carcinogenesis and in response to EoE inflammation. Here, we employed scRNA-Seq to compare the impact of these two conditions upon the esophageal epithelial landscape. In mice with EoE, we detect accumulation of 4 suprabasal populations and depletion of 4 basal cell populations. These findings are consistent with studies demonstrating impaired squamous differentiation in human subjects with EoE (43). However, it must be noted that while EoE patients often feature basal cell hyperplasia, a histological finding in which basal cells occupy >25% of esophageal epithelial cell height (44), our findings in mice with EoE show a depletion of basal populations. Here, we have defined basal, suprabasal, and superficial cells using a combination of unbiased bioinformatics-based clustering along with mapping of putative markers of basal and superficial cells onto the identified cell clusters. Although basal cell hyperplasia is assumed to occur via expansion of the basal cell compartment, presumably via a proliferative response, it is possible that basal cell hyperplasia as seen in EoE may instead result from an accumulation of suprabasal cells that fail to differentiate. In an independent scRNA-Seq dataset, we recently validated ATP1B3 as a marker of suprabasal cells in murine esophageal epithelium (40). As such, it will be of interest to determine how EoE inflammation impacts the expression of ATP1B3. A recent human scRNA-Seq study identified an increase in proliferating suprabasal cells, but not proliferating basal cells, in the epithelium of active EoE subjects (45). Identification of proliferating cells in the noted human EoE dataset was based on expression of the proliferation marker *TOP2A* (45). Although we did not evaluate *Top2a*, analysis of cell cycle genes in our dataset does not support proliferation across the EoE-enriched suprabasal populations and pathway analysis in these clusters predicted an association with senescence pathway. It is possible that studies in human subjects may have captured EoE across a broader spectrum of disease activity as compared to our murine studies which comprised littermates from the inbred C57B6 strain treated with an identical protocol to induce EoE. Future studies using time course experiments in murine EoE cohorts may resolve the apparent discrepancies with regard to cell proliferation in humans and mice with EoE inflammation. In contrast to mice with EoE, mice with ESCC displayed evidence of basal cell hyperplasia with accumulation of 4 basal populations, including basal population 4 in which S-phase genes were enriched. Pathway analysis further identified an association between cell proliferation pathways, including EIF2 signaling, regulation of eIF4 p70S6 Kinase, and mTOR, in ESCC-enriched cell populations. Collectively, our findings support a model wherein basal hyperplasia in EoE results from stalled squamous differentiation and accumulation of suprabasal cells while basal cell hyperplasia in ESCC results from accumulation of highly proliferative basal cells (**Figure 9**).

**Figure 9.**
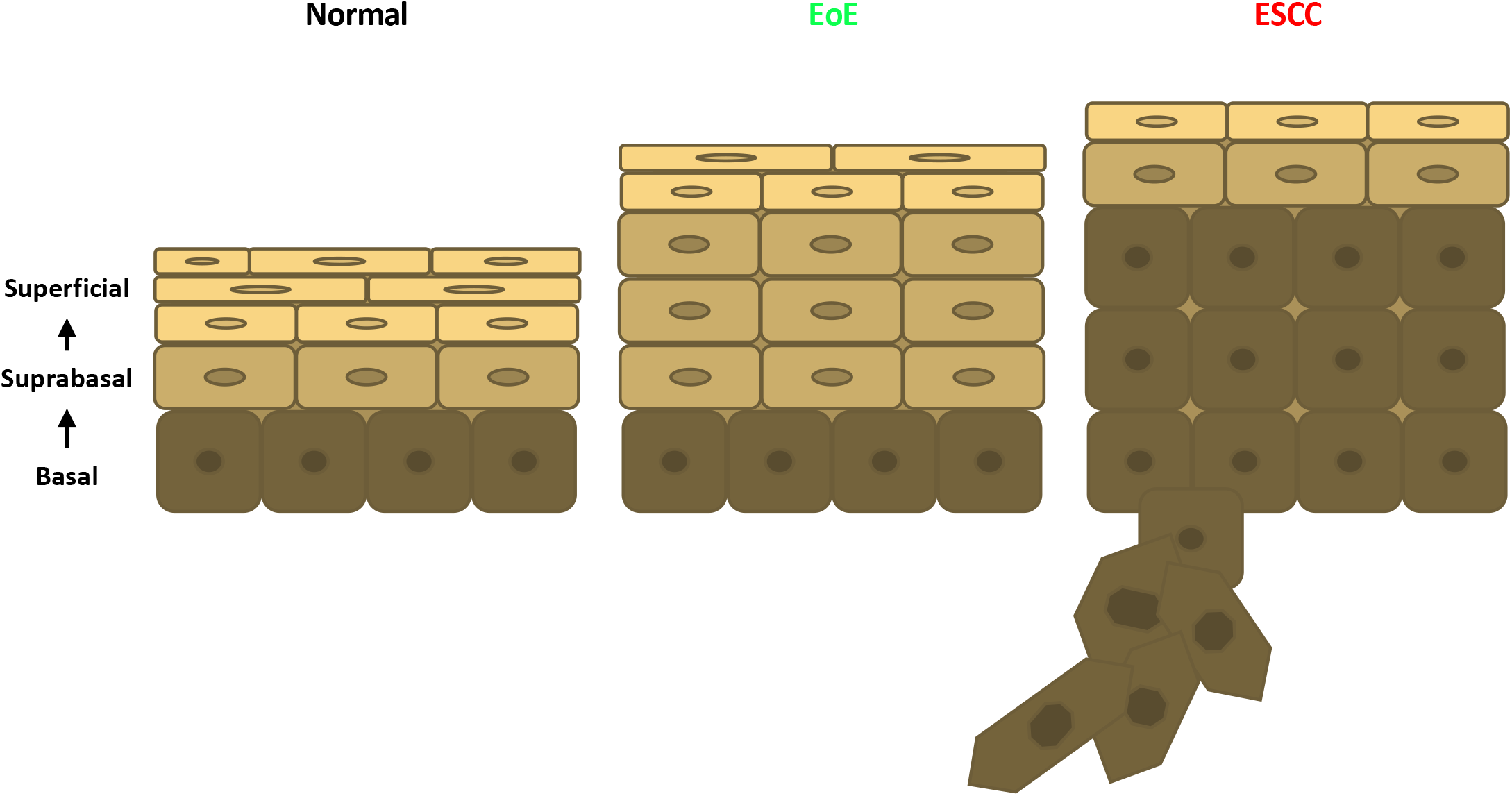
Schematic models of esophageal epithelial cell development between normal, EoE, and ESCC conditions. **(A)** Graphical representation of cell differentiation between conditions. In EoE, an expansion of suprabasal cells is present in the epithelium. In ESCC, an expansion of basal cells is present in the epithelium and encroaches into the stroma.

Our scRNA-Seq-based findings in mice with EoE or ESCC coupled with decreased tumor burden in mice with ESCC and EoE as compared to those with ESCC only, supports the premise that exposure to EoE inflammation may alter esophageal epithelial biology in a manner that effectively limits esophageal carcinogenesis. By pairing murine models of EoE and ESCC we provide the first functional assessment of the relationship between these two disease entities and report that exposure to EoE inflammation limits esophageal carcinogenesis. These findings agree with epidemiological studies in EoE patients which have failed to identify esophageal malignancy in EoE patients (19, 20, 46). The relationship between allergy and cancer remains elusive. While a number of epidemiological studies report a negative association between allergic inflammation and cancer (20–28, 46), others report positive or null associations (47, 48). A major limitation of such population-based studies is the inherent heterogeneity of human cohorts with regard to genetics and environmental exposures. A strength of the current study is the use of murine models of esophageal food allergy and cancer that allow for the study of the interaction of these conditions while minimizing effects of such confounding variables. Conversely, the current study lacks scRNA-Seq data in mice with EoE and ESCC in combination. Such investigations are currently underway and will examine the mutational burden and epigenetic landscape in mice with EoE and ESCC alone and in combination to determine if exposure to EoE inflammation may limit activation of oncogenes and/or enhance activation of tumor suppressors.

Although the current study examined effects of EoE and ESCC on esophageal epithelium, it is likely that immune-mediated pathways contribute to EoE-mediated suppression of esophageal carcinogenesis. As EoE is clinically characterized by the presence of eosinophils, it is tempting to speculate that these cells may be integral to EoE-mediated suppression of esophageal carcinogenesis. In ESCC, high eosinophil counts are associated with favorable patient outcomes (36, 37, 49, 50). Melanoma patients with high eosinophil counts have also been demonstrated to have prolonged survival following immunotherapy (51). In preclinical models, IL-33 delays metastatic progression of peritoneal cancer via induction of an allergic tumor microenvironment in which eosinophils, CD4+ T cells, and macrophages contribute to anti-tumoral effects (52). Moreover, while direct eosinophil-mediated cytotoxicity has been demonstrated *in vitro*, eosinophil-dependent inhibition of tumor initiation *in vivo* occurs by IL-33-mediated effects upon CD8+ T cells in the tumor microenvironment (53). Eosinophil recruitment of CD8+ T cells has also been shown to facilitate tumor rejection *in vivo* (54). Mast cells have also been implicated in EoE pathogenesis (55–57); however, the relationship of these cell types to cancer remains elusive. The presence of mast cell is associated with improved prognosis in colorectal and breast cancers (58, 59); however, mast cells have also been linked to tumor promotion via induction of resistance to anti-PD-1 therapy in melanoma and IL-33-elicited macrophage mobilization in murine models of gastric cancer (60, 61). In ESCC, mast cells have been implicated in tumor progression via angiogenesis (62). Here, we utilized epithelium-enriched esophageal tissue fractions for scRNA-Seq which precluded evaluation of the impact of EoE and ESCC on inflammatory cells. In our ongoing studies of mice with EoE and ESCC both alone and in combination, whole esophagi will be subjected to scRNA-Seq identify immune cells that may contribute to EoE-mediated suppression of esophageal carcinogenesis.

In sum, this investigation unveils marked differences in esophageal epithelial cell remodeling occurring in EoE as compared to ESCC. While esophageal epithelium of mice with ESCC features accumulation of basal cells, including those with proliferative capacity, esophageal epithelium of mice with EoE features accumulation of suprabasal cells that are largely in G0/G1-phase of the cell cycle and express a gene profile that is associated with senescence. *In vivo* studies combining murine models of EoE and ESCC further demonstrate that exposure to EoE inflammation limits ESCC carcinogenesis. Taken together, our findings raise the possibility that exposure to allergic inflammation may inhibit carcinoma development in the esophagus by pushing esophageal epithelial cells toward a state of stalled differentiation in which they lack the proliferative potential that is needed for carcinogenesis. Should this notion be validated, it may illuminate novel approaches based on epithelial cell fate reprogramming for cancer prevention and/or therapy.

## 5 Life Science Identifiers

Not applicable

## 6 Conflict of Interest

The authors declare that the research was conducted in the absence of any commercial or financial relationships that could be construed as a potential conflict of interest.

## 7 Author Contributions

ADF: Data curation, formal analysis and writing—original draft; ALK: formal and statistical analysis; MFK: collection of data; AK: writing—review and editing; JLJ: writing—review and editing; AM: collection of the data; YT: sequencing; AKS: formal analysis; KAW: conceptualization, initiation of the study, supervision and writing—reviewing and editing. All authors contributed to the article and approved the submitted version.

## 8 Funding

The following grant supported this work: R01DK121159 (KAW), R21CA256465 (KAW), R01DK121159-S1 (JLJ), T32GM142606 (ADF; PIs: Xavier Graña, Jonathan Soboloff, Temple University), P30CA006927 (YT, AKS; PI: Jonathan Chernoff, Fox Chase Cancer Center).

## 9 Acknowledgments

We thank Fox Chase Cancer Center Histopathology Facility for technical support.

## 10 Contribution to the Field Statement

Although emerging evidence supports the premise that allergy-associated alterations in immune signaling suppress cancer development and progression, our understanding of how epithelial barrier surfaces exposed to allergic inflammation may affect carcinogenesis is sparse. Here, we performed single cell RNA-sequencing in murine models of esophageal cancer (i.e. esophageal squamous cell carcinoma) and food allergy (i.e. eosinophilic esophagitis) to examine the impact of these two conditions upon the esophageal epithelial cell landscape. Although basal cell hyperplasia has been reported in both conditions, our study indicates that mice with esophageal cancer display accumulation of proliferative basal cells while mice with esophageal food allergy exhibit accumulation of suprabasal cells that are predicted to be post-mitotic and have enrichment of senescence-associated genes. In addition, *in vivo* data indicates an ability of food allergy to limit tumorigenesis using these murine models. In sum, this study provides novel insight into the functional relationship between EoE and ESCC and may have future implications for leveraging allergic inflammation-associated alterations in epithelial biology to prevent and/or treat esophageal cancer.

## 11 Data Availability Statement

The authors declare that all data supporting the findings of this study are available within the article and its supplementary information files or from the corresponding author upon reasonable request.

## Notes

https://www.ncbi.nlm.nih.gov/geo/query/acc.cgi?acc=GSE218118

### Competing Interest Statement

The authors have declared no competing interest.

## References

1. Takahashi Y, Fukui T, Kishimoto M, Suzuki R, Mitsuyama T, Sumimoto K, et al. Phosphorylation of Smad2/3 at the specific linker threonine residue indicates slow-cycling esophageal stem-like cells before re-entry to the cell cycle. Dis Esophagus. 2016;29(2):107–15.

2. Messier B, Leblond CP. Cell proliferation and migration as revealed by radioautography after injection of thymidine-H3 into male rats and mice. Am J Anat. 1960;106:247–85.

3. Al Yassin TM, Toner PG. Fine structure of squamous epitheilum and submucosal glands of human oesophagus. J Anat. 1977;123(Pt 3):705–21.

4. Hanawa M, Suzuki S, Dobashi Y, Yamane T, Kono K, Enomoto N, et al. EGFR protein overexpression and gene amplification in squamous cell carcinomas of the esophagus. Int J Cancer. 2006;118(5):1173–80.

5. Song Y, Li L, Ou Y, Gao Z, Li E, Li X, et al. Identification of genomic alterations in oesophageal squamous cell cancer. Nature. 2014;509(7498):91–5.

6. Network CGAR, University AWGA, Agency BC, Hospital BaWs, Institute B, University B, et al. Integrated genomic characterization of oesophageal carcinoma. Nature. 2017;541(7636):169–75.

7. Hollstein MC, Metcalf RA, Welsh JA, Montesano R, Harris CC. Frequent mutation of the p53 gene in human esophageal cancer. Proc Natl Acad Sci U S A. 1990;87(24):9958–61.

8. Agrawal N, Jiao Y, Bettegowda C, Hutfless SM, Wang Y, David S, et al. Comparative genomic analysis of esophageal adenocarcinoma and squamous cell carcinoma. Cancer Discov. 2012;2(10):899–905.

9. Rustgi AK, El-Serag HB. Esophageal carcinoma. N Engl J Med. 2014;371(26):2499–509.

10. Cook MB, Corley DA, Murray LJ, Liao LM, Kamangar F, Ye W, et al. Gastroesophageal reflux in relation to adenocarcinomas of the esophagus: a pooled analysis from the Barrett’s and Esophageal Adenocarcinoma Consortium (BEACON). PLoS One. 2014;9(7):e103508.

11. Orlando RC. Pathophysiology of gastroesophageal reflux disease. J Clin Gastroenterol. 2008;42(5):584–8.

12. Souza RF, Huo X, Mittal V, Schuler CM, Carmack SW, Zhang HY, et al. Gastroesophageal reflux might cause esophagitis through a cytokine-mediated mechanism rather than caustic acid injury. Gastroenterology. 2009;137(5):1776–84.

13. Merves J, Muir A, Modayur Chandramouleeswaran P, Cianferoni A, Wang ML, Spergel JM. Eosinophilic esophagitis. Ann Allergy Asthma Immunol. 2014;112(5):397–403.

14. Katzka DA, Ravi K, Geno DM, Smyrk TC, Iyer PG, Alexander JA, et al. Endoscopic Mucosal Impedance Measurements Correlate With Eosinophilia and Dilation of Intercellular Spaces in Patients With Eosinophilic Esophagitis. Clin Gastroenterol Hepatol. 2015;13(7):1242–8.e1.

15. Capocelli KE, Fernando SD, Menard-Katcher C, Furuta GT, Masterson JC, Wartchow EP. Ultrastructural features of eosinophilic oesophagitis: impact of treatment on desmosomes. J Clin Pathol. 2015;68(1):51–6.

16. Collins MH. Histopathologic features of eosinophilic esophagitis. Gastrointest Endosc Clin N Am. 2008;18(1):59–71; viii-ix.

17. Whelan KA, Godwin BC, Wilkins B, Elci OU, Benitez A, DeMarshall M, et al. Persistent Basal Cell Hyperplasia Is Associated With Clinical and Endoscopic Findings in Patients With Histologically Inactive Eosinophilic Esophagitis. Clin Gastroenterol Hepatol. 2020;18(7):1475–82.e1.

18. Vicario M, Blanchard C, Stringer KF, Collins MH, Mingler MK, Ahrens A, et al. Local B cells and IgE production in the oesophageal mucosa in eosinophilic oesophagitis. Gut. 2010;59(1):12–20.

19. Lipka S, Keshishian J, Boyce HW, Estores D, Richter JE. The natural history of steroid-naïve eosinophilic esophagitis in adults treated with endoscopic dilation and proton pump inhibitor therapy over a mean duration of nearly 14 years. Gastrointest Endosc. 2014;80(4):592–8.

20. Syed A, Maradey-Romero C, Fass R. The relationship between eosinophilic esophagitis and esophageal cancer. Dis Esophagus. 2017;30(7):1–5.

21. Cui Y, Hill AW. Atopy and Specific Cancer Sites: a Review of Epidemiological Studies. Clin Rev Allergy Immunol. 2016;51(3):338–52.

22. Cockcroft DW, Klein GJ, Donevan RE, Copland GM. Is there a negative correlation between malignancy and respiratory atopy? Ann Allergy. 1979;43(6):345–7.

23. Grulich AE, Vajdic CM, Kaldor JM, Hughes AM, Kricker A, Fritschi L, et al. Birth order, atopy, and risk of non-Hodgkin lymphoma. J Natl Cancer Inst. 2005;97(8):587–94.

24. La Vecchia C, D’Avanzo B, Negri E, Franceschi S. History of selected diseases and the risk of colorectal cancer. Eur J Cancer. 1991;27(5):582–6.

25. La Vecchia C, Negri E, D’Avanzo B, Boyle P, Franceschi S. Medical history and primary liver cancer. Cancer Res. 1990;50(19):6274–7.

26. Bueno de Mesquita HB, Maisonneuve P, Moerman CJ, Walker AM. Aspects of medical history and exocrine carcinoma of the pancreas: a population-based case-control study in The Netherlands. Int J Cancer. 1992;52(1):17–23.

27. Negri E, Bosetti C, La Vecchia C, Levi F, Tomei F, Franceschi S. Allergy and other selected diseases and risk of colorectal cancer. Eur J Cancer. 1999;35(13):1838–41.

28. Wang H, Diepgen TL. Atopic dermatitis and cancer risk. Br J Dermatol. 2006;154(2):205–10.

29. Wang H, Rothenbacher D, Löw M, Stegmaier C, Brenner H, Diepgen TL. Atopic diseases, immunoglobulin E and risk of cancer of the prostate, breast, lung and colorectum. Int J Cancer. 2006;119(3):695–701.

30. Zacharia BE, Zacharia B, Sherman P. Atopy, helminths, and cancer. Med Hypotheses. 2003;60(1):1–5.

31. Josephs DH, Spicer JF, Corrigan CJ, Gould HJ, Karagiannis SN. Epidemiological associations of allergy, IgE and cancer. Clin Exp Allergy. 2013;43(10):1110–23.

32. McCraw AJ, Chauhan J, Bax HJ, Stavraka C, Osborn G, Grandits M, et al. Insights from IgE Immune Surveillance in Allergy and Cancer for Anti-Tumour IgE Treatments. Cancers (Basel). 2021;13(17).

33. Chatterjee J, Sanapala S, Cobb O, Bewley A, Goldstein AK, Cordell E, et al. Asthma reduces glioma formation by T cell decorin-mediated inhibition of microglia. Nat Commun. 2021;12(1):7122.

34. Poli A, Oudin A, Muller A, Salvato I, Scafidi A, Hunewald O, et al. Allergic airway inflammation delays glioblastoma progression and reinvigorates systemic and local immunity in mice. Allergy. 2022.

35. Noti M, Wojno ED, Kim BS, Siracusa MC, Giacomin PR, Nair MG, et al. Thymic stromal lymphopoietin-elicited basophil responses promote eosinophilic esophagitis. Nat Med. 2013;19(8):1005–13.

36. Naganuma S, Whelan KA, Natsuizaka M, Kagawa S, Kinugasa H, Chang S, et al. Notch receptor inhibition reveals the importance of cyclin D1 and Wnt signaling in invasive esophageal squamous cell carcinoma. Am J Cancer Res. 2012;2(4):459–75.

37. Tang XH, Knudsen B, Bemis D, Tickoo S, Gudas LJ. Oral cavity and esophageal carcinogenesis modeled in carcinogen-treated mice. Clin Cancer Res. 2004;10(1 Pt 1):301–13.

38. Whelan KA, Merves JF, Giroux V, Tanaka K, Guo A, Chandramouleeswaran PM, et al. Autophagy mediates epithelial cytoprotection in eosinophilic oesophagitis. Gut. 2017;66(7):1197–207.

39. Li M, Hener P, Zhang Z, Kato S, Metzger D, Chambon P. Topical vitamin D3 and low-calcemic analogs induce thymic stromal lymphopoietin in mouse keratinocytes and trigger an atopic dermatitis. Proc Natl Acad Sci U S A. 2006;103(31):11736–41.

40. Kabir MF, Karami AL, Cruz-Acuña R, Klochkova A, Saxena R, Mu A, et al. Single cell transcriptomic analysis reveals cellular diversity of murine esophageal epithelium. Nat Commun. 2022;13(1):2167.

41. Nennstiel S, Schlag C. Treatment of eosinophlic esophagitis with swallowed topical corticosteroids. World J Gastroenterol. 2020;26(36):5395–407.

42. Rochman M, Xie YM, Mack L, Caldwell JM, Klingler AM, Osswald GA, et al. Broad transcriptional response of the human esophageal epithelium to proton pump inhibitors. J Allergy Clin Immunol. 2021;147(5):1924–35.

43. Muir AB, Wang JX, Nakagawa H. Epithelial-stromal crosstalk and fibrosis in eosinophilic esophagitis. J Gastroenterol. 2019;54(1):10–8.

44. Steiner SJ, Kernek KM, Fitzgerald JF. Severity of basal cell hyperplasia differs in reflux versus eosinophilic esophagitis. J Pediatr Gastroenterol Nutr. 2006;42(5):506–9.

45. Rochman M, Wen T, Kotliar M, Dexheimer PJ, Ben-Baruch Morgenstern N, Caldwell JM, et al. Single-cell RNA-Seq of human esophageal epithelium in homeostasis and allergic inflammation. JCI Insight. 2022;7(11).

46. Straumann A, Spichtin HP, Grize L, Bucher KA, Beglinger C, Simon HU. Natural history of primary eosinophilic esophagitis: a follow-up of 30 adult patients for up to 11.5 years. Gastroenterology. 2003;125(6):1660–9.

47. Vaengebjerg S, Skov L, Egeberg A, Loft ND. Prevalence, Incidence, and Risk of Cancer in Patients With Psoriasis and Psoriatic Arthritis: A Systematic Review and Meta-analysis. JAMA Dermatol. 2020;156(4):421–9.

48. Kantor ED, Hsu M, Du M, Signorello LB. Allergies and Asthma in Relation to Cancer Risk. Cancer Epidemiol Biomarkers Prev. 2019;28(8):1395–403.

49. Natsuizaka M, Whelan KA, Kagawa S, Tanaka K, Giroux V, Chandramouleeswaran PM, et al. Interplay between Notch1 and Notch3 promotes EMT and tumor initiation in squamous cell carcinoma. Nat Commun. 2017;8(1):1758.

50. Yang X, Wang L, Du H, Lin B, Yi J, Wen X, et al. Prognostic impact of eosinophils in peripheral blood and tumor site in patients with esophageal squamous cell carcinoma treated with concurrent chemoradiotherapy. Medicine (Baltimore). 2021;100(3):e24328.

51. Platzer B, Elpek KG, Cremasco V, Baker K, Stout MM, Schultz C, et al. IgE/FcεRI-Mediated Antigen Cross-Presentation by Dendritic Cells Enhances Anti-Tumor Immune Responses. Cell Rep. 2015;10(9):1487–95.

52. Perales-Puchalt A, Svoronos N, Villarreal DO, Zankharia U, Reuschel E, Wojtak K, et al. IL-33 delays metastatic peritoneal cancer progression inducing an allergic microenvironment. Oncoimmunology. 2019;8(1):e1515058.

53. Lucarini V, Ziccheddu G, Macchia I, La Sorsa V, Peschiaroli F, Buccione C, et al. IL-33 restricts tumor growth and inhibits pulmonary metastasis in melanoma-bearing mice through eosinophils. Oncoimmunology. 2017;6(6):e1317420.

54. Carretero R, Sektioglu IM, Garbi N, Salgado OC, Beckhove P, Hämmerling GJ. Eosinophils orchestrate cancer rejection by normalizing tumor vessels and enhancing infiltration of CD8(+) T cells. Nat Immunol. 2015;16(6):609–17.

55. Niranjan R, Mavi P, Rayapudi M, Dynda S, Mishra A. Pathogenic role of mast cells in experimental eosinophilic esophagitis. Am J Physiol Gastrointest Liver Physiol. 2013;304(12):G1087–94.

56. Nelson M, Zhang X, Pan Z, Spechler SJ, Souza RF. Mast cell effects on esophageal smooth muscle and their potential role in eosinophilic esophagitis and achalasia. Am J Physiol Gastrointest Liver Physiol. 2021;320(3):G319–G27.

57. Dellon ES, Chen X, Miller CR, Fritchie KJ, Rubinas TC, Woosley JT, et al. Tryptase staining of mast cells may differentiate eosinophilic esophagitis from gastroesophageal reflux disease. Am J Gastroenterol. 2011;106(2):264–71.

58. Nielsen HJ, Hansen U, Christensen IJ, Reimert CM, Brünner N, Moesgaard F. Independent prognostic value of eosinophil and mast cell infiltration in colorectal cancer tissue. J Pathol. 1999;189(4):487–95.

59. Rajput AB, Turbin DA, Cheang MC, Voduc DK, Leung S, Gelmon KA, et al. Stromal mast cells in invasive breast cancer are a marker of favourable prognosis: a study of 4,444 cases. Breast Cancer Res Treat. 2008;107(2):249–57.

60. Somasundaram R, Connelly T, Choi R, Choi H, Samarkina A, Li L, et al. Tumor-infiltrating mast cells are associated with resistance to anti-PD-1 therapy. Nat Commun. 2021;12(1):346.

61. Eissmann MF, Dijkstra C, Jarnicki A, Phesse T, Brunnberg J, Poh AR, et al. IL-33-mediated mast cell activation promotes gastric cancer through macrophage mobilization. Nat Commun. 2019;10(1):2735.

62. Elpek GO, Gelen T, Aksoy NH, Erdoğan A, Dertsiz L, Demircan A, et al. The prognostic relevance of angiogenesis and mast cells in squamous cell carcinoma of the oesophagus. J Clin Pathol. 2001;54(12):940–4.

